# Origin of elevational replacements in a clade of nearly flightless birds – most diversity in tropical mountains accumulates via secondary contact following allopatric speciation

**DOI:** 10.1101/606558

**Authors:** Carlos Daniel Cadena, Laura N. Céspedes

## Abstract

Tropical mountains are biodiversity hotspots. In particular, mountains in the Neotropics exhibit remarkable beta diversity reflecting species turnover along elevational gradients. Elevational replacements of species have been known since early surveys of the tropics, but data on how such replacements arise are scarce, limiting our understanding of mechanisms underlying patterns of diversity. We employed a phylogenetic framework to evaluate hypotheses accounting for the origin of elevational replacements in the genus *Scytalopus* (Rhinocryptidae), a speciose clade of passerine birds with limited dispersal abilities occurring broadly in the Neotropical montane region. We found that species of *Scytalopus* have relatively narrow elevational ranges, closely related species resemble each other in elevational distributions, and most species replacing each other along elevational gradients are distantly related to each other. Although we cannot reject the hypothesis that a few elevational replacements may reflect parapatric speciation along mountain slopes, we conclude that speciation in *Scytalopus* occurs predominantly in allopatry within elevational zones, with most elevational replacements resulting from secondary contact of formerly allopatric lineages. Our study suggests that accumulation of species diversity in montane environments reflects colonization processes as opposed to *in situ* divergence even in dispersal-limited animals.

## Introduction

Species turnover along elevational gradients is a salient pattern in tropical biogeography. Ever since pioneering work by Francisco José de Caldas and Alexander von Humboldt on plant geography, naturalists have noticed that many species occur over narrow ranges of elevation and replace each other along mountain slopes (Nieto, 2006; von Humboldt & Bonpland, 2009). Elevational replacements of closely related species are prevalent in the tropics (Terborgh, 1971; Diamond, 1973; Wake & Lynch, 1976), where organisms likely have narrower physiological tolerances than in temperate zones (Janzen, 1967; McCain, 2009). Marked changes in species assemblages with elevation (e.g. of plants, invertebrates and vertebrates; Patterson *et al.*, 1998; Kessler, 2001; Jankowski *et al.*, 2013b; García-Robledo *et al.*, 2016; Gill *et al.*, 2016; Badgley *et al.*, 2018) thus result in tropical mountains being hotspots of beta diversity (Melo *et al.*, 2009; Fjeldså *et al.*, 2012). Therefore, knowledge about evolutionary and ecological mechanisms involved in the origin of elevational replacements is central to understanding major patterns in the distribution of life (Janzen, 1967; Huey, 1978).

Abutting species distributions along elevational gradients may reflect either (1) parapatric ecological speciation leading to divergence of a formerly widespread species into two or more daughter species with restricted ranges, or (2) secondary contact following range expansions of species originating in allopatry (Endler, 1982; Hua, 2016). Although one should exercise caution when making inferences about the geographic context of speciation based on current geographic distributions (Losos & Glor, 2003), these alternative hypotheses are, in principle, amenable to testing by means of phylogenetic analyses: parapatric divergence predicts that species replacing each other with elevation are sister to each other, whereas secondary contact predicts they are not (Patton & Smith, 1992; Moritz *et al.*, 2000). The few studies testing these predictions on animals indicate that most speciation in the montane Neotropics occurs in allopatry and that species replacing each other along elevational gradients are not each other’s closest relatives (Patton & Smith, 1992; Cadena *et al.*, 2012; Caro *et al.*, 2013). Therefore, elevational replacements may primarily reflect secondary contact but more studies are necessary to confirm this pattern.

As evidenced by some of the first large-scale surveys of the geographic and ecological distributions of species (Chapman, 1917; Todd & Carriker, 1922; Chapman, 1926), most birds living in montane areas of the Neotropics have narrow elevational ranges (Jankowski *et al.*, 2013a; but see Gadek *et al.*, 2018 for exceptions). For example, median ranges of species across their geographic distributions span only ca. 1100 m in three families of Neotropical birds (Figure 1; see also Graves, 1988). Narrow elevational ranges of individual species are often coupled with segregation with ecologically similar species along mountain slopes. For instance, a series of landmark studies in the Peruvian Andes documented multiple cases of pairs of congeneric species of birds replacing each other with elevation as well as scenarios where up to 4-5 congeners occur successively along mountain slopes with minimum overlap (Terborgh, 1971; Terborgh & Weske, 1975; Terborgh, 1977). Understanding ecological and physiological mechanisms maintaining patterns of elevational segregation in tropical birds has been the focus of multiple studies (reviewed by Jankowski *et al.*, 2013a), yet analyses of the origins of elevational replacements remain scarce (Cadena, 2007; Freeman, 2015; Cadena *et al.*, 2019a). In particular, we are unaware of studies attempting to test alternative hypotheses accounting for the origin of elevational replacements in a diverse clade of widespread organisms in the context of a robust phylogeny.

**Figure 1.**
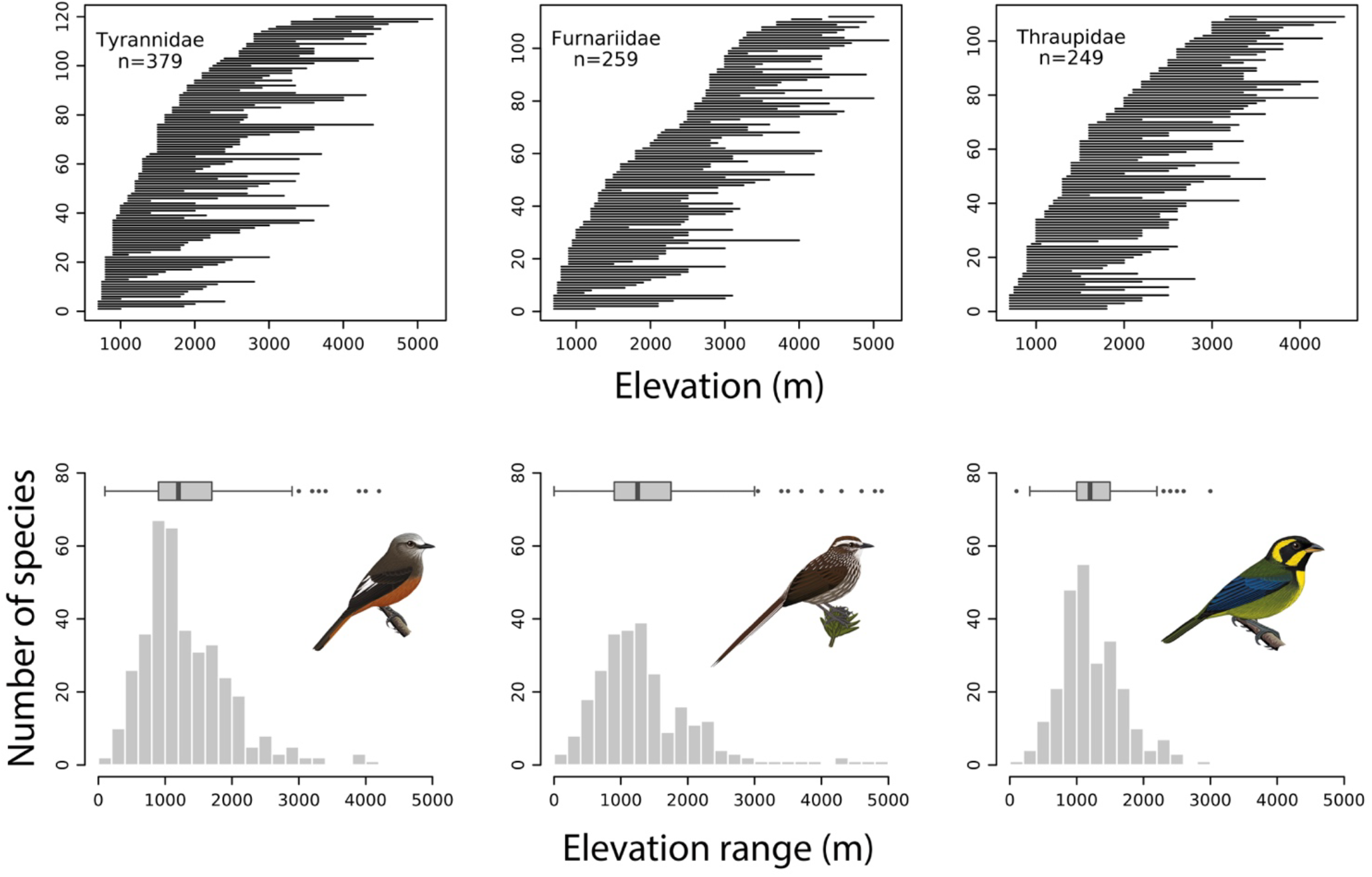
Restricted elevational distributions of species in three diverse families of Neotropical birds: ovenbirds (Furnariidae), tyrant flycatchers (Tyrannidae), and tanagers (Thraupidae). The panels in the top row depict the range of elevations occupied by species with lower elevational limits at or above 700 m ordered from lowest to highest along the vertical axis, showing turnover of species with elevation. Panels in the bottom row summarize the data above with histograms and boxplots, indicating that most species inhabit only a fraction of the elevational ranges existing on mountains like the Andes; across the three families, median elevational ranges span only ca. 1100 m and very few species have ranges greater than 2500 m. Data are from Parker *et al*. (1996) and illustrations from Ayerbe-Quiñones (Ayerbe-Quiñones, 2018).

Tapaculos in the genus *Scytalopus* (Rhinocryptidae) are small passerine birds with poor dispersal abilities ranging broadly in the Neotropical montane region. Except for the Pantepui, species in the genus occupy all major mountains in the Neotropics from Costa Rica to Patagonia, reaching lowland areas and foothills in southern South America and eastern Brazil. *Scytalopus* are mouse-like birds which forage by walking or hopping on or near the ground in dense vegetation; they are unable to engage in long, powered flights because they have small and rounded wings and unfused clavicles (Figure 2; Krabbe & Schulenberg, 2003; Maurício *et al.*, 2008). *Scytalopus* avoid highly lit open areas and are rarely found far from vegetation cover except in barren high-elevation environments. Local diversity of *Scytalopus* is typically low, yet multiple species may be found in different habitats in a given landscape, being prime examples of patterns of elevational replacement (Krabbe & Schulenberg, 1997, 2003). For example, on a single morning walking trails upslope in forests along the Cerro de Montezuma on the Pacific-facing slope of the Andes of Colombia, birdwatchers may successively encounter *S. chocoensis*, *S. alvarezlopezi, S. vicinior*, *S. spillmanni*, and *S. latrans*; if they visit paramos at higher elevations in the region, they may also find *S. canus* (Stiles *et al.*, 2017; see below). Elevational replacements in the genus are typically sharp. Species seldom co-occur at the same elevations and when they do so, they often segregate by habitat. Current taxonomy recognizes 44 species of *Scytalopus*, but this is most certainly an underestimate given marked genetic structure within species, geographic variation in vocalizations, and the potential for discovery of new taxa in unexplored regions (Cadena *et al.*, 2019b).

**Figure 2.**
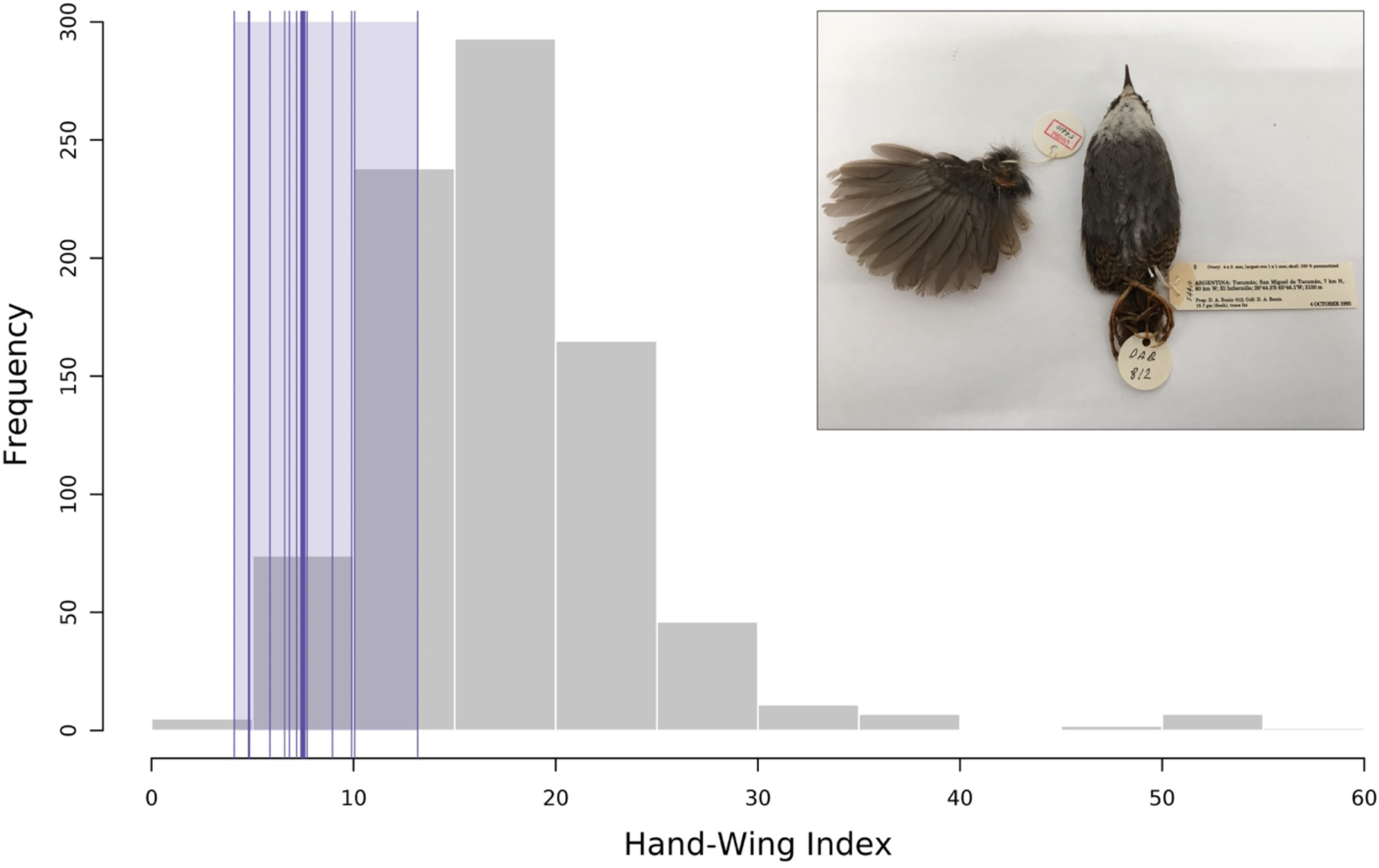
The frequency distribution of the hand-wing index, a proxy for dispersal abilities in birds, shows that *Scytalopus* tapaculos exhibit reduced potential for flighted dispersal relative to other birds. The dimensionless hand-wing index describes wing size and shape as a function of wing length (measured from the carpal joint to the longest primary feather) and secondary length (measured from the carpal joint to the tip of the first secondary feather), with larger values indicating greater dispersal abilities (Claramunt *et al.*, 2012). The histogram in grey depicts the distribution of the hand-wing index across a sample of 851 species of New World passerine birds (measurements from Claramunt *et al.*, 2012; P. Montoya, G. Bravo, and E. Tenorio, unpubl. data). Data for *Scytalopus* are in purple, showing the median (vertical thick bar) and the range (shaded area) of the hand-wing index across 12 species (thin bars are mean values per species). The inset illustrates the small and rounded wing of a specimen of *S. superciliaris* from Argentina housed at the Burke Museum of Natural History and Culture at the University of Washington (photograph by Cooper French).

Elevational replacements in *Scytalopus* are somewhat paradoxical given both main hypotheses posed to account for such patterns. First, given the morphological and life-history traits making these birds poor dispersers one would expect that long-distance dispersal or range expansions would hardly bring multiple species together in a given slope, especially because mountain regions are separated by geographic barriers associated with population isolation and diversification in other birds (Hazzi *et al.*, 2018). Alternatively, given what we know about the geographic context of speciation in birds in general (Phillimore *et al.*, 2008; Price, 2008) and in the Andes in particular (García-Moreno & Fjeldså, 2000; Caro *et al.*, 2013), parapatric divergence of multiple species along elevational gradients in several geographic regions would also appear unlikely. A preliminary analysis of diversification in *Scytalopus* focusing on few taxa from eastern Ecuador and Peru suggested secondary contact is a more likely explanation for elevational replacements (Arctander & Fjeldså, 1994; Roy *et al.*, 1997). However, because the phylogeny employed for analyses may have been problematic (Arctander, 1995), taxonomic and geographic coverage were limited, and the substantial improvement of our understanding of geographic ranges, species limits, and diversity in *Scytalopus* (Krabbe & Schulenberg, 1997; Cadena *et al.*, 2019b), revisiting these questions is warranted. We used a comprehensive molecular phylogeny and data on geographic and elevational distributions of species to describe the mode of speciation and thereby examine the origin of elevational replacements in *Scytalopus*.

## Methods

A recent study reconstructed molecular phylogenies of *Scytalopus* employing sequences of mitochondrial DNA (mtDNA), 80 nuclear exons, and > 1800 regions flanking ultraconserved elements in the nuclear genome (Cadena *et al.*, 2019b). We used the mtDNA ND2 data set from that study, which had denser taxonomic and geographic sampling, as the basis for our analyses; phylogenies inferred with this data set were largely congruent with those based on nuclear data for nodes relevant to analyses we present here. We inferred a gene tree using Beast 2.4.8 (Bouckaert *et al.*, 2014) for an alignment of 90 sequences, including 83 *Scytalopus* and seven outgroups. We applied a relaxed uncorrelated clock (mean=0.0125, SD=0.15; Smith & Klicka, 2010) and a Yule speciation tree prior. We ran chains for 100 million generations and discarded the initial 50% as burn-in.

We were able to gather elevational distribution data for 57 taxa of *Scytalopus*; these represent all named species as well as several distinct populations, which may be undescribed species given genetic divergence, vocal variation, or geographic distributions (see Cadena *et al.*, 2019b). We obtained information on elevational ranges from taxonomic descriptions and other papers (Whitney, 1994; Krabbe & Schulenberg, 1997; Cuervo *et al.*, 2005; Krabbe *et al.*, 2005; Maurício, 2005; Bornschein *et al.*, 2007; Donegan & Avendaño-C., 2008; Vasconcelos *et al.*, 2008; Krabbe & Cadena, 2010; Whitney *et al.*, 2010; Donegan *et al.*, 2013; Hosner *et al.*, 2013; Maurício *et al.*, 2014; Avendaño *et al.*, 2015; Avendaño & Donegan, 2015; Stiles *et al.*, 2017), regional handbooks (Fjeldså & Krabbe, 1990; Ridgely & Greenfield, 2001; Hilty, 2003; Restall *et al.*, 2007; Schulenberg *et al.*, 2007; Herzog *et al.*, 2016; Ayerbe-Quiñones, 2018), and the Handbook of the Birds of the World (del Hoyo *et al.*, 2018). Additionally, fo several taxa we defined elevational distributions based on expert knowledge (A M. Cuervo, N. K. Krabbe, D. F. Lane, T. S. Schulenberg, and V. Piacentini, unpubl. data); this was particularly useful for unnamed populations differing phenotypically or genetically from others. We performed all analyses described below using all 57 taxa and separately for a data set including only 47 of them which represent all currently recognized species (American Ornithologists’ Union 1998; Stiles *et al.*, 2017; Remsen *et al.*, 2018) plus three unnamed species from the Andes of Peru (an unnamed form referred to *S. altirostris* mentioned by Cadena *et al.*, 2019b was not considered). To conduct analyses, we trimmed the phylogeny constructed with the complete data set (i.e., 90 terminals) to include only the 57 or 47 taxa considered. The elevation ranges for species in the 47-taxon data set encompassing more than one taxon in the 57-taxon data set correspond to the combined ranges of these taxa (e.g. the range of *S. rodriguezi* in the 47-taxon data set is 1700-2300 m, i.e. the combined range of forms *rodriguezi and yariguiorum*).

To describe changes in elevational distributions over the history of *Scytalopus*, we mapped the midpoint of the elevation range of each taxon in the phylogeny using the *contMap* function of *phytools*, which estimates character states at nodes and along branches using a maximum-likelihood approach (Revell, 2012). Additionally, we calculated Pagel’s λ (Pagel, 1999) as a measure of phylogenetic signal of midpoint elevation and accounted for phylogenetic uncertainty by calculating this statistic across the final 1,001 trees in the posterior distribution using *phytools*. Using likelihood-ratio tests, we also tested whether Pagel’s λ in each tree was significantly different from 0 (i.e. no phylogenetic signal) and 1 (i.e. the value expected under pure Brownian motion). Phylogenetic signal is often interpreted in terms of the degree of conservatism or lability of a trait (e.g. Blomberg *et al.*, 2003), but inferences about factors underlying patterns should be cautious because different evolutionary processes may produce similar values of phylogenetic signal (Revell *et al.*, 2008). Nonetheless, high phylogenetic signal can be interpreted as a strong tendency of closely related species to resemble each other in a given trait (Revell *et al.*, 2008). Therefore, high phylogenetic signal in elevational ranges could indicate close resemblance between close relatives in such ranges. In addition to examining the midpoint of elevational ranges of species, we also conducted analyses based on minimum and maximum elevation, obtaining qualitatively similar results.

To evaluate whether sister taxa have similar or contrasting elevational distributions as predicted by allopatric and parapatric speciation, respectively (Patton & Smith, 1992), we identified sister taxa across the 1,001 final trees in the posterior distribution. We then calculated the elevational overlap of pairs of sister taxa by dividing the amount of overlap by the elevational range of the taxon with the narrowest range (Kozak & Wiens, 2007; Cadena *et al.*, 2012). A value of 1 indicates that either ranges are exactly the same or that the narrower range is entirely contained in the wider range; a value of 0 indicates that elevational distributions do not overlap. The number of pairs of sister taxa employed for analyses varied across trees in the posterior from 18 to 23 (median = 20.3, total across trees = 51 pairs) in the 57-tip data set, and from 12 to 18 (median = 15.1, total across trees = 52 pairs) in the 47-tip data set. These analyses were restricted to sister taxa representing terminal branches in the phylogeny (i.e., we did not employ ancestral state reconstructions to compare elevational ranges in cases when a species representing a long terminal branch was sister to a clade formed by ≥ 2 species).

Finally, we graphically examined the phylogenetic affinities of species of *Scytalopus* replacing each other along elevational gradients in four regions of South America to examine whether such replacements more likely reflect secondary contact or parapatric speciation along mountain slopes. These regions were: (A) the Sierra Nevada de Santa Marta in northern Colombia, (B) the Pacific slope of the Western Cordillera of Colombia, (C) Zamora-Chinchipe Province on the Amazonian slope of the Andes of Ecuador, and (D) the Río Satipo Valley in Junín Department, eastern Andean slope of Peru. All analyses were conducted and figures plotted in the R programming environment (R Core Team 2018).

## Results

*Scytalopus* tapaculos jointly occupy a wide range of elevations in the Neotropics, from sea level up to 4,600 m in the high Andes (Table 1, Figure 3). Elevational ranges vary substantially among species, from very narrow (200 m) to quite broad (3,500 m), yet most species occupy only a relatively small fraction of the elevational gradients in which they occur: mean ranges were 1079 m (SD= 625 m) in the 57-taxa data set and 1166 m (SD= 641 m) in the 47-taxa data set. Given the geographic setting where most species of *Scytalopus* occur, where mountains reach very high altitudes and habitats for birds may extend over several thousand meters (e.g., Figure 1), the elevational ranges of species are generally rather narrow. Overall, taxa within main clades of *Scytalopus* have roughly similar elevational distributions. For example, most species in the Southern Andean clade (Figure 3, clade B) occur exclusively at high elevations; exceptions include *S. fuscus*, found in lowlands, and *S*. *magellanicus*, a temperate-zone species with the widest range in the genus (0 to 3,500 m). Species from the tropical Andes and Central America (Figure 3, clade C) show wide variation in elevational distributions, but species within subclades in the region tend to have similar ranges. For example, species in clades D, G and I all occur at high elevations in the tropical Andes except for *S. femoralis*, *S. micropterus* and *S. caracae*, which inhabit lower elevations than their close relatives (Figure 3). All taxa in clades E, F and H occur at mid elevations, with some ranging to lower elevations (Figure 3). All species from Brazil (Figure 3, clade A) occur at low to mid elevations, an unsurprising pattern given that mountains reach much lower elevations in that region than in the Andes.

**Table 1.**
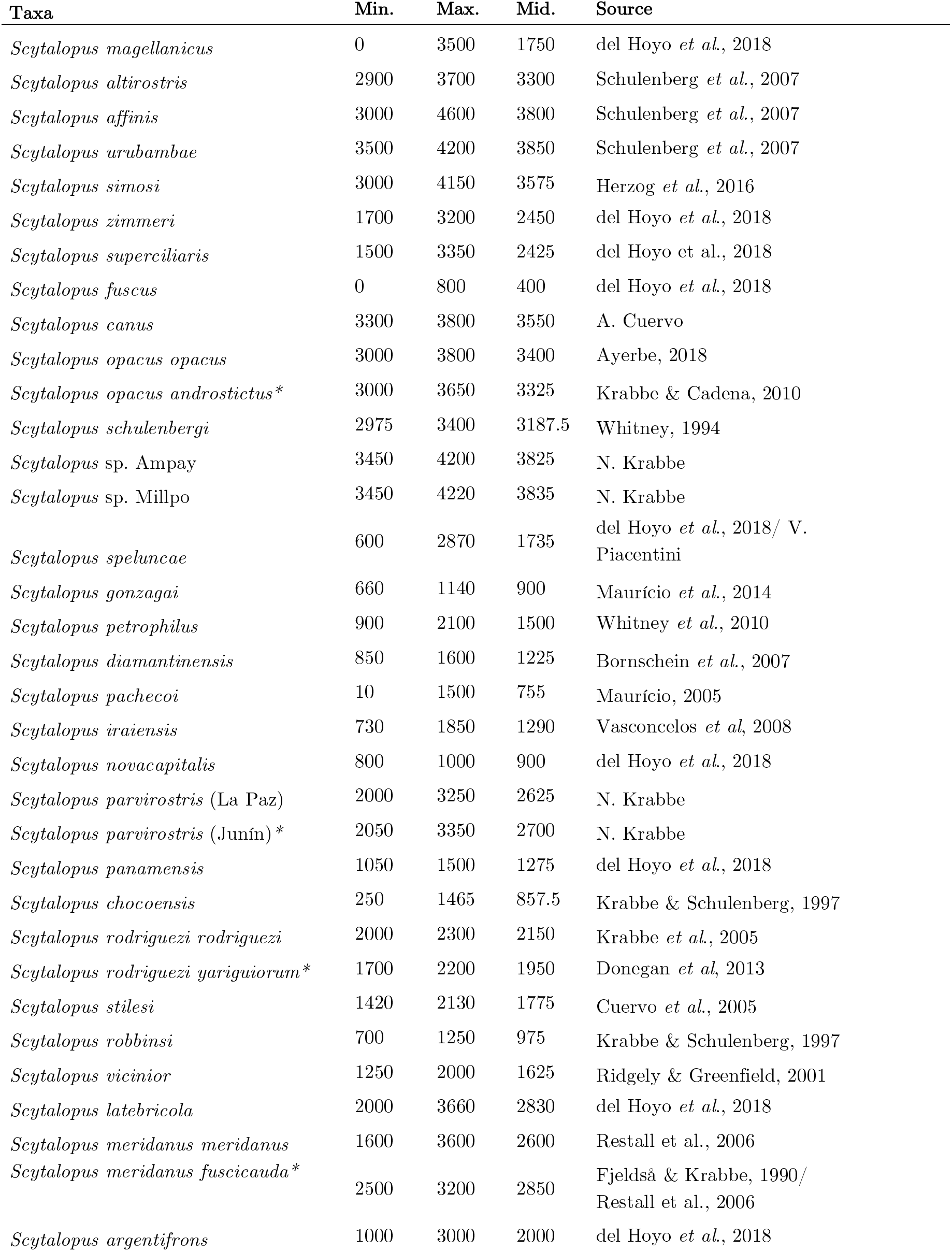
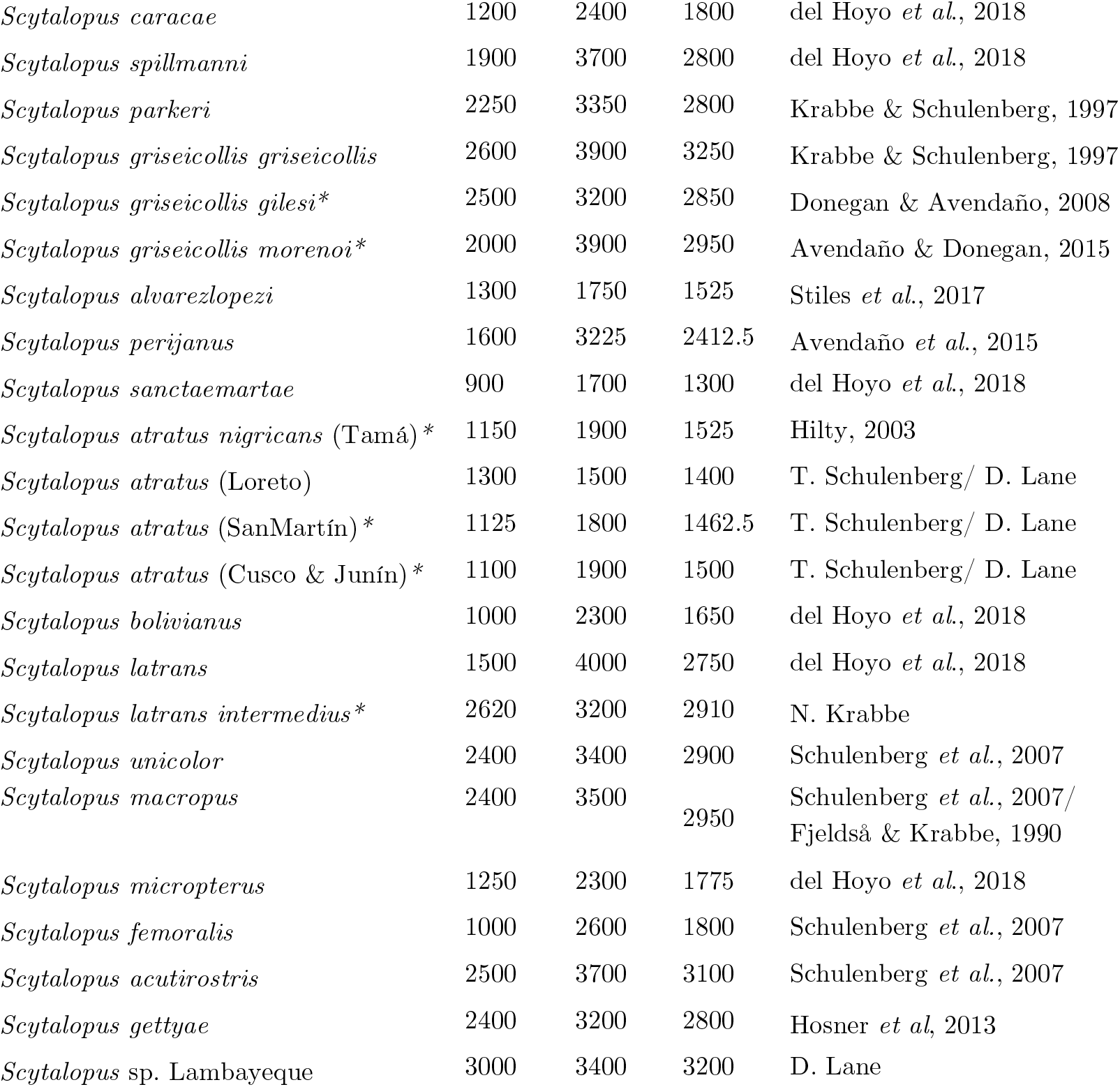
Elevational distributions of *Scytalopus* tapaculos considered in analyses. For each taxon, we provide the minimum, maximum and midpoint of the elevation range (in m), as well as sources for these data. Taxa with asterisks were those excluded from analyses involving only the 47 species recognized (or soon to be recognized) by taxonomists (see text).

**Figure 3.**
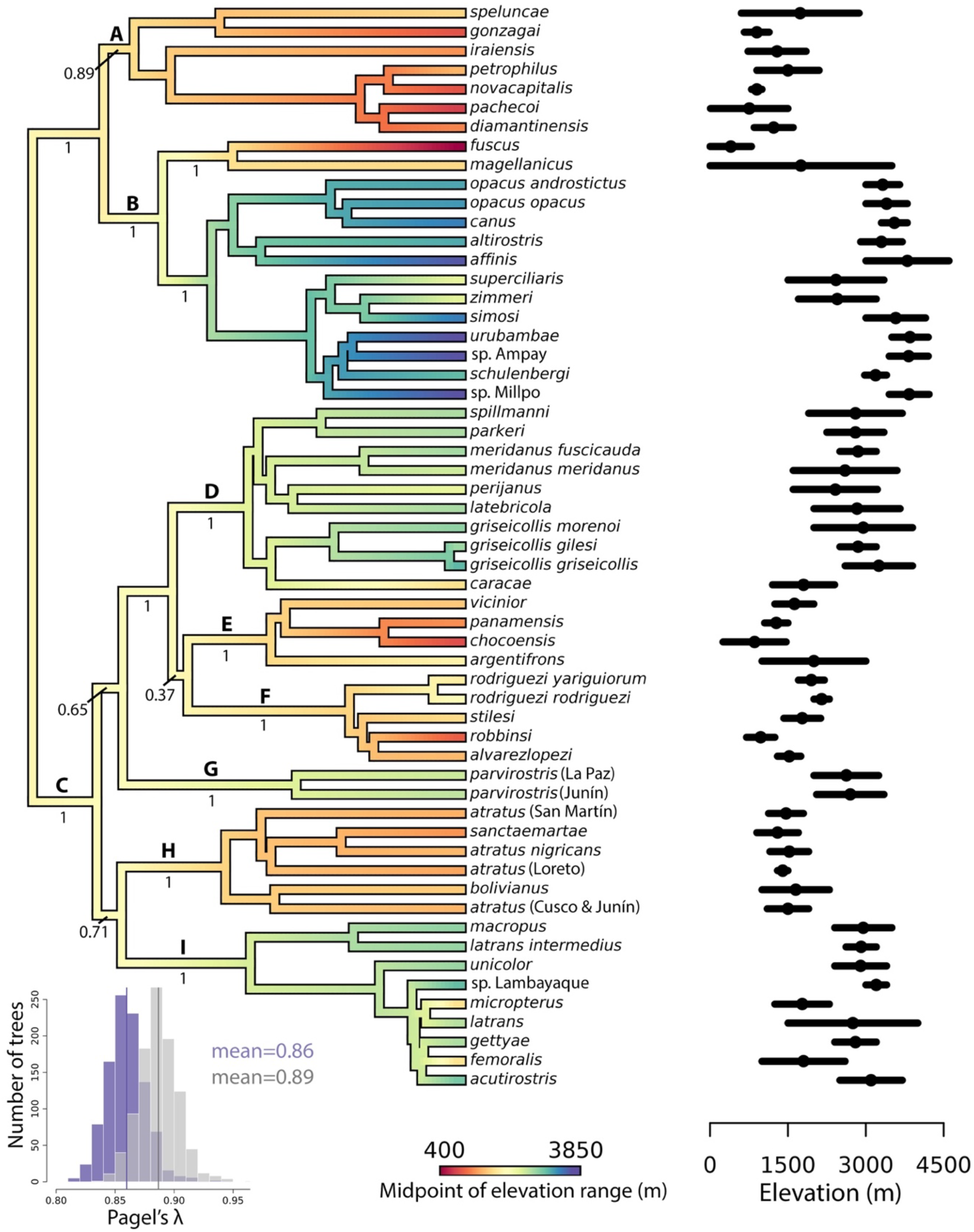
Elevational ranges of species are broadly similar within main clades of *Scytalopus* tapaculos and there is relatively high phylogenetic signal in midpoint of the elevational range. The phylogeny is the maximum clade credibility tree with midpoint of elevation mapped on branches using maximum-likelihood; posterior probabilities are shown only for major clades. The elevation range and midpoint elevation of each taxon are shown in front of each tip. The histogram in the bottom left depicts the distribution of phylogenetic signal in midpoint elevation measured by Pagel’s λ estimated across 1,001 posterior trees (purple for the 47-taxa data set, gray for the 57-taxa data set).

Phylogenetic signal for the midpoint of the elevational range of species measured by Pagel’s λ, which differed significantly from 0 across all trees and for both data sets (p<0.001), was relatively high (mean=0.89 for the 57-taxon data set and 0.86 for the 47-taxon data set; Figure 3). However, in most trees (77% in the 57-taxon data set and 64% in the 47-taxon data set) phylogenetic signal was also significantly different from 1 (p<0.05), implying that elevational ranges are more divergent than expected given evolution under pure Brownian motion. These results indicate that closely related *Scytalopus* tend to strongly resemble each other in the midpoint of their elevation ranges, but differences among species cannot be fully accounted for by time since their divergence. The minimum and maximum elevations in the ranges of species also had significant phylogenetic signal; estimates of Pagel’s λ for maximum elevation were very similar to those we obtained for midpoint elevation, while those for minimum elevation were slightly lower yet qualitatively similar (data not shown).

Most sister taxa in *Scytalopus* have similar elevational distributions (Figure 4). Mean overlap in elevational ranges between sister taxa was slightly higher in the 57-taxon data set (pooled mean across the 1,001 trees=0.76, SD= 0.33) than in the 47-taxon data set (mean=0.70, SD=0.38). Regardless of the data set employed for analyses, half or more pairs of sister taxa overlapped substantially in elevational ranges (overlap >0.8) while less than a quarter of pairs of sister taxa had distinct elevational ranges (overlap <0.2; Figure 4). Furthermore, the majority of sister taxa identified across the 1,001 trees (10 pairs out of 11, or 12 out of 13 depending on the data set) showing overlap <0.2 do not occur on the same gradient, i.e. they are allopatric. These results suggest speciation in *Scytalopus* occurs predominantly within elevational zones and not in parapatry along mountain slopes. The only possible exception to this pattern is divergence between *S. acutirostris* and *S. femoralis*, which may or may not be sister species, but are close relatives replacing each other along the Amazonian slope of the Central Andes of Peru (see below). We note that our analyses included data for at most 57 taxa yet the true number of species in *Scytalopus* is likely higher. Because most of the lineages which we did not consider are closely allied to other lineages with similar elevational ranges (e.g. groups within *S. atratus* or *S. parvirostris;* Cadena *et al*. 2019b) we believe that if there is any bias in our results it would be in the direction of underestimating the true overlap in elevational ranges of close relatives. In other words, greater taxonomic coverage would likely reinforce our conclusion that sister taxa have similar elevational distributions.

**Figure 4.**
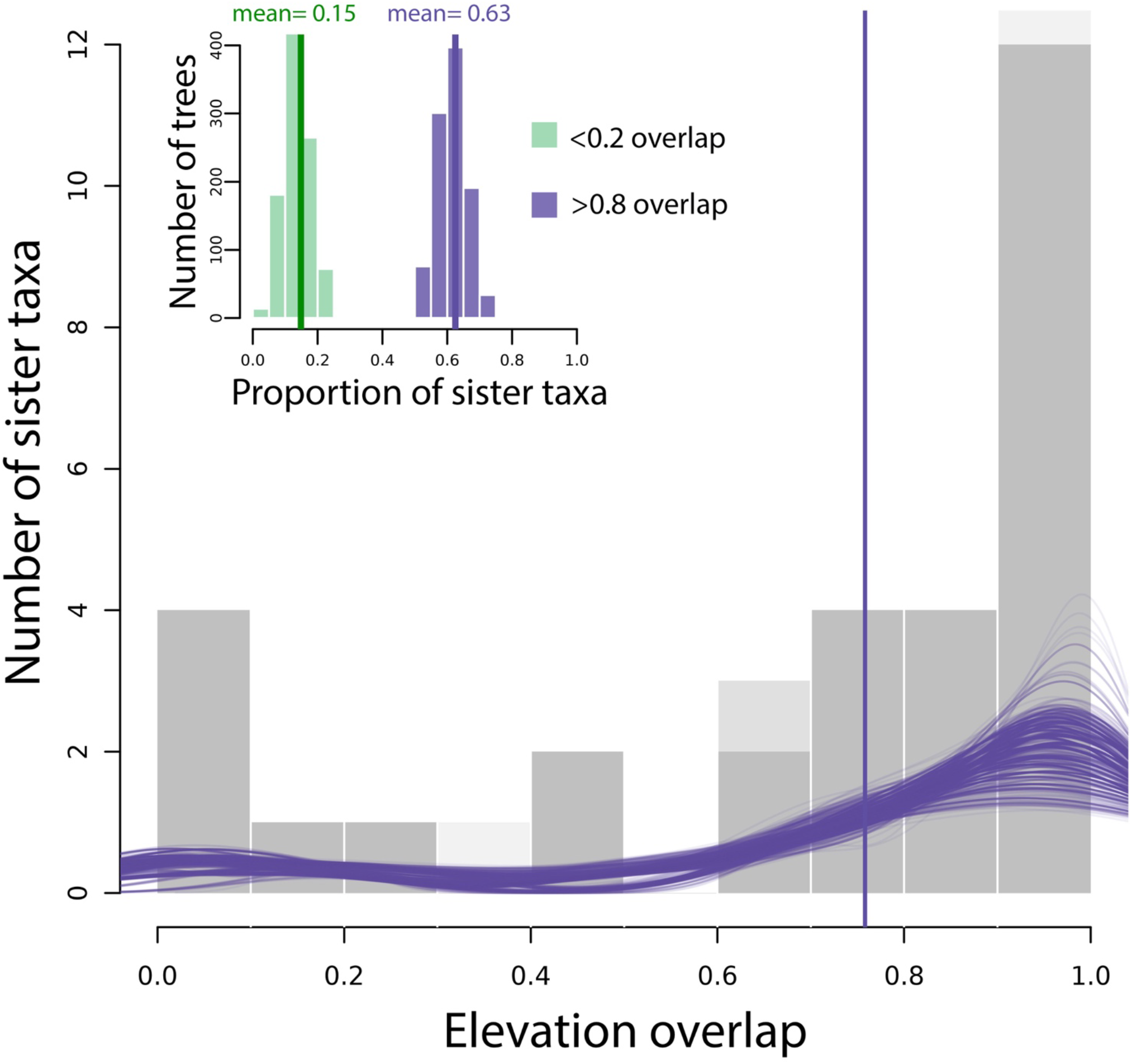
Most sister species of *Scytalopus* have similar elevational distributions. The main figure shows an overlay of 1,001 histograms, each corresponding to a phylogenetic tree in the posterior, showing the distribution of values of elevational overlap between sister taxa (0 indicates no overlap, 1 indicates complete overlap of elevational ranges); purple lines are density plots and the vertical line signals the pooled mean across trees. Histograms in the inset show the distribution of the proportion of sister taxa in the 57-taxon data set with little (<0.2) and high overlap (>0.8) in elevational distributions across the 1,001 trees. We obtained qualitatively similar results with the 47-taxon data set (see text).

The majority of *Scytalopus* replacing each other along the elevation gradients we examined are distant relatives (Figure 5). The two species endemic to the Sierra Nevada de Santa Marta belong to distinct clades last sharing a common ancestor ca. 8 million years ago (clades D and H in Figure 2; Figure 5). Likewise, the four species occurring in forests on the Pacific slope of the Western Andes of Colombia belong to four different clades (clades D, E, F and I in Figure 3), whereas a fifth species (*S. canus*) occurring in páramo habitats above treeline belongs to yet another clade (clade B). The species found above and below *S. alvarezlopezi* in the region (*S. vicinior* and *S. chocoensis*, respectively; Figure 5) belong to clade E, but they are not sister to each other. The five species replacing each other along the Amazonian slope of the Andes in Zamora-Chinchipe, Ecuador, belong to four distinct clades (B, D, H and I in Figure 3; Figure 5). Two of the species with parapatric distributions in this gradient (*S. latrans* and *S. micropterus*) are closely related, but the posterior probability of the hypothesis that they are sisters is only 0.43. The most recent common ancestor of species occurring in elevational gradients in western Colombia, in eastern Ecuador, and in eastern Peru is the most recent common ancestor of all *Scytalopus*, which existed ca. 9.8 million years ago (crown age 7.8-12.3 m.a. highest posterior density; Figure 5). Taken together, the above data indicate that most species in the genus replacing each other along elevational gradients in South America did not evolve in parapatry *in situ*, but rather met in each gradient following divergence in allopatry.

**Figure 5.**
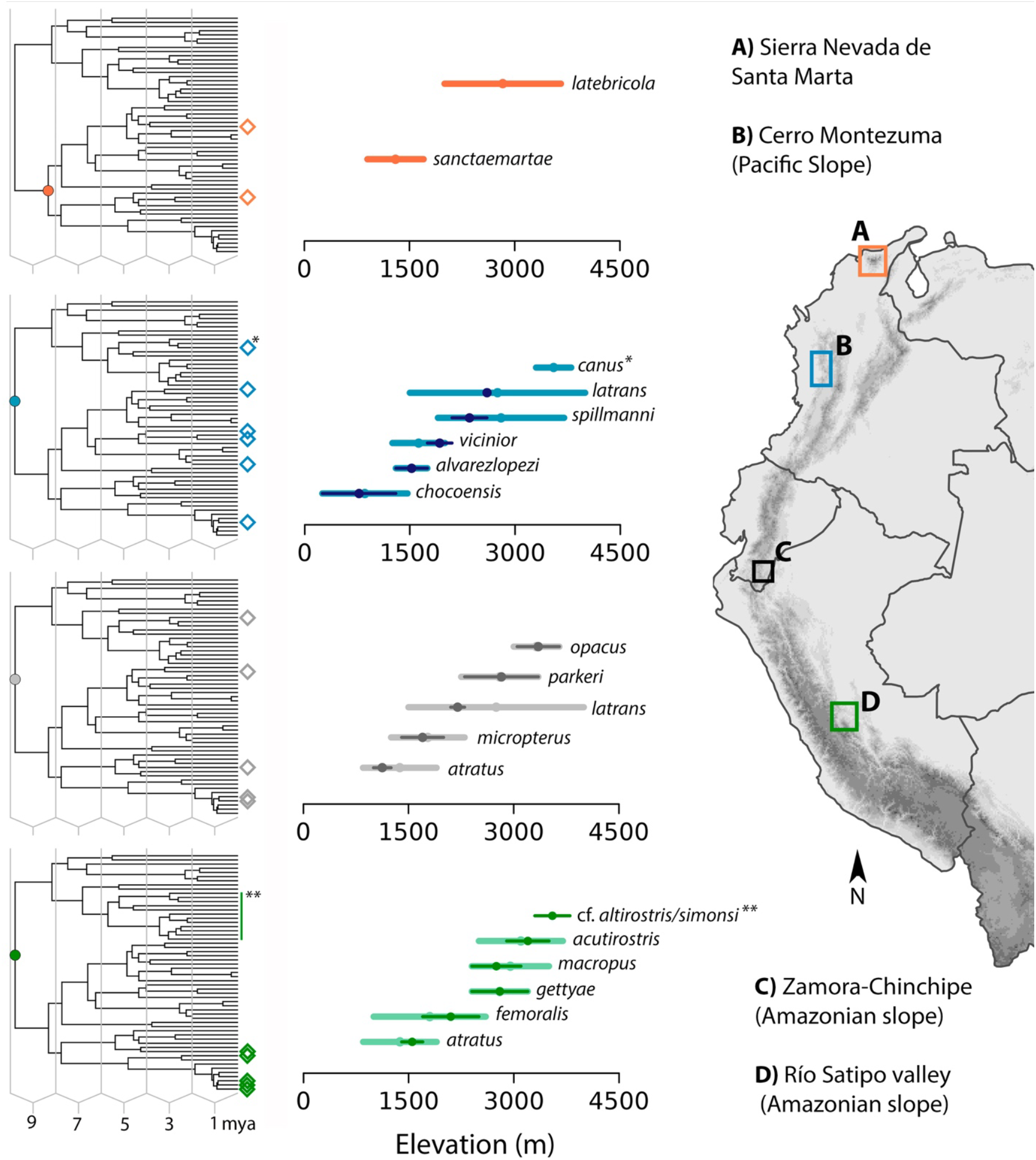
*Scytalopus* species replacing each other with elevation in four regions of South America are often, but not always, distantly related. Rhombuses at the tips of the phylogeny highlight species that replace each other along each elevational gradient, and the dot denotes the most recent common ancestor of these species. Horizontal bars depict the elevational range of species, where the lighter color represents the elevation range for the species across its distribution and the darker color the elevational range reported for each specific gradient (B: Stiles *et al*., 2017; C: Krabbe & Schulenberg, 1997; D: Hosner *et al*., 2013); a single bar is shown for the species from the Sierra Nevada de Santa Marta because both are endemic and for *S. canus* because the species does not occur in the specific gradient we highlight (i.e., Cerro Montezuma). The range of *S. latrans* in the Cerro de Montezuma is shown as a dot because it is only known from the very highest elevations in the area. Although the taxonomic identity of the species occurring at the highest elevations in the Satipo Valley gradient is uncertain, it very likely belongs to the clade depicted with the vertical line on the tree (Hosner *et al.*, 2013).

However, the possibility of parapatric speciation remains plausible for some of the species replacing each other with elevation on the eastern slope of the Andes. In contrast to patterns observed in other regions, four of the six species replacing each other with elevation the Río Satipo Valley in Junín, Peru, belong to clade I (Figure 3; Figure 5). Moreover, three of these species (*S. femoralis, S. gettyae* and *S. acutirostris*) belong to a group nested within clade I including several closely allied taxa with shallow divergence in mtDNA (Figure 3; Figure 5). The remaining two species found in this gradient belong to clade B (an unidentified taxon; Hosner *et al*., 2013) and clade H (*S. atratus*). Unpublished evidence indicates a seventh species (*S.* aff. *parvirostris*) belonging to clade G also occurs in the Río Satipo Valley (N. Krabbe, pers. comm.) but we did not include it in Figure 5 because information on its elevational range in the area is lacking.

## Discussion

Many species of birds and other organisms have restricted elevational distributions, particularly in the tropics. This results in biodiversity patterns observable globally (e.g., tropical mountains are hotspots of species turnover in space because species replace each other with elevation; Fjeldså *et al.*, 2012) and regionally (e.g., diversity may peak at mid elevations or decline with elevation in a given mountain; Quintero & Jetz, 2018). We probed into evolutionary processes resulting in replacements of species along mountain slopes by examining the elevational ranges of species in the context of a phylogeny of *Scytalopus* tapaculos, a speciose clade of Neotropical montane birds in which the elevational replacement of species is commonplace. We found that (1) elevational ranges of species of *Scytalopus* are relatively narrow given the broad elevational gradients of the mountains where they live, (2) closely related species in the genus usually resemble each other in elevational distributions, (3) most pairs of sister taxa have largely overlapping elevational ranges, and (4) species coexisting regionally with elevational segregation on mountains are very often –but not always– distantly related to each other. Assuming that current distributional ranges are informative about the geographic context of speciation (Barraclough & Vogler, 2000), our study thus suggests that speciation in *Scytalopus* occurs predominantly in allopatry within elevational zones, and that elevational replacements typically result from secondary contact of formerly allopatric species rather than from primary divergence in parapatry (see also Arctander & Fjeldså, 1994). However, the latter possibility cannot be entirely excluded for some species and regions.

Our analyses of phylogenetic signal in elevational ranges and of overlap in elevational ranges between sister taxa treat such ranges as if they were phenotypic attributes amenable to evolution. Although elevational ranges are not organismal traits, we consider them emergent properties of populations which do reflect heritable phenotypes allowing organisms to persist over a given range of environments, justifying our study of the evolution of such ranges in a phylogenetic framework (Cadena, 2007). We acknowledge, however, that because elevation *per se* is not a factor directly influencing organisms, one needs to exercise caution when comparing elevational distributions among species living in regions differing in the way in which elevation covaries with conditions and resources to which organisms directly respond (Cadena & Loiselle, 2007). In this regard, we point out that some factors likely limiting organisms such as partial O_2_ pressure do vary consistently with elevation regardless of geographic context, but others like temperature do not. However, because closely related species of *Scytalopus* typically occur in the same geographic region (i.e. at similar latitudes; Cadena *et al.*, 2019b), we believe it is generally sensible to assume that similar elevational distributions reflect similar ecology and underlying phenotypic traits.

In agreement with our results, previous work on birds (García-Moreno & Fjeldså, 2000; Caro *et al.*, 2013) and other animals (Patton & Smith, 1992; Lynch, 1999) indicates that species replacing each other with elevation in the Neotropics are often not sister to each other. Furthermore, sister species in several vertebrate clades overlap considerably in their elevational ranges in Neotropical mountains (Cadena *et al.*, 2012), suggesting that speciation occurs most often in allopatry within elevational zones and thus that elevational replacements result predominantly from secondary contact (but see Kozak & Wiens, 2007). Work on this topic in other tropical regions has been more limited, yet evidence from Africa (Fuchs *et al.*, 2011) and southeast Asia (Moyle *et al.*, 2017) also indicates secondary contact is the most likely explanation for elevational replacements (but see Bryja *et al.*, 2018; Eldridge *et al.*, 2018). Likewise, assembly of biotas in other mountain systems often results more from colonization by lineages from other regions than from diversification within mountains (Johansson *et al.*, 2007; Merckx *et al.*, 2015). Therefore, understanding the processes influencing the dynamics of geographic ranges which lead to secondary sympatry following divergence in allopatry is central to establishing how diversity accumulates in montane regions.

Those unfamiliar with *Scytalopus* tapaculos might be unsurprised by our finding that species originating in distinct mountains may come together into regional sympatry with elevational segregation in a given mountain. After all, tapaculos are birds and birds fly around. We, however, find this result quite striking because, unlike many birds, *Scytalopus* are notably poor dispersers (Krabbe & Schulenberg, 2003). Birds in the genus walk and hop much more than they fly, have tiny and rounded wings which preclude long-distance powered flight, and have even lost fused clavicles, one of the most exquisite putative adaptations of birds in general to their flighted life style. The behavior of *Scytalopus* also makes them highly reluctant to disperse: most species very rarely venture far from forest cover, having been described as agoraphobic (Krabbe & Schulenberg, 1997) or photophobic (Sick, 1993). Even during storms in the high Andes, tapaculos tend to stay put: rather than moving downslope to avoid inclement weather, individuals maintain their territories and forage in tunnel systems under the snow (Fjeldså, 1991).

How did such undispersive birds manage to get around, colonizing an isolated mountain system like the Sierra Nevada de Santa Marta twice or the two slopes of the northern Andes multiple times? Tapaculos are not alone in achieving such feats. Phylogeographic analyses of *Henicorhina* wood-wrens (Troglodytidae), another group of poorly dispersive birds, also revealed that elevational replacements result from secondary contact of formerly allopatric lineages (Caro *et al.*, 2013; Cadena *et al.*, 2019a). An explanation for the apparent paradox of poor dispersers repeatedly coming into contact from disjunct areas, even in highly isolated mountains, is that individual birds did not disperse over large distances crossing barriers now appearing unsurmountable. Rather, populations likely tracked the dynamics of their favored environments, gradually expanding their geographic distributions in concert with climatic change. During cool periods in Earth history, montane environments in tropical mountains were displaced downslope, which increased opportunities for formerly isolated areas to become connected by vegetation; in turn, isolation among such areas likely increased during warmer periods when vegetation zones retreated upslope (Hooghiemstra & Van der Hammen, 2004; Bush *et al.*, 2011). Repeated cycles of disconnection and connection of montane areas (Ramírez-Barahona & Eguiarte, 2013) may thus have spurred cycles of allopatric speciation and subsequent secondary contact, thereby enabling the regional accumulation of diversity (Roy *et al.*, 1997). Owing to the expected narrow thermal tolerance of tropical montane organisms (Janzen, 1967), one would expect this mechanism of divergence and accumulation of diversity in mountains to be especially prevalent in the tropics (Ghalambor *et al.*, 2006; Kozak & Wiens, 2007; Cadena *et al.*, 2012). An alternative, nonexclusive explanation for cycles of allopatric speciation followed by establishment of secondary sympatry, is that species may go through phases of expansion and contraction of their geographic ranges even in the absence of marked changes in the physical environment (Cadena *et al.*, 2019a). This may occur owing to evolution of phenotypic traits influencing dispersal (Hosner *et al.*, 2017), or to changes in ecological specialization and interactions with other species (Ricklefs, 2010).

Although adaptation to divergent selective pressures along gradients of elevation may seem like a prime precursor to the origin of new species (e.g. Funk *et al.*, 2016; Hua, 2016), parapatric speciation along mountain slopes appears to be rare. In contrast to data discussed above, several studies do suggest that species replacing each other along elevational gradients may be closely related (Bates & Zink, 1994; Hall, 2005; DuBay & Witt, 2012), yet evidence that these replacements do not reflect separate colonization events of elevation belts or lowland-highland vicariance resulting from uplift processes (Brumfield & Edwards, 2007; Ribas *et al.*, 2007; Santos *et al.*, 2009) is lacking. To our knowledge, the clearest example of parapatric speciation in mountains involves sister species in the plant genus *Senecio* occurring on Mount Etna, Italy, which differ strikingly in ecology and phenotype despite experiencing extensive gene flow (Chapman *et al.*, 2013; Osborne *et al.*, 2013; Chapman *et al.*, 2016). A promising additional case is that of *Syma* kingfishers in New Guinea, in which two distinct species co-occuring with elevational segregation have experienced gene flow yet maintain divergence in regions of the genome likely involved in adaptation and, presumably, mate choice (Linck *et al.*, 2019). In contrast to the *Senecio* and *Syma* examples, avian taxa replacing each other with elevation in Neotropical mountains seldom show evidence of gene flow, with the only documented cases of interbreeding between elevational replacements in the region we are aware of being those of *Anairetes* tit-tyrants in Peru (Dubay & Witt, 2014), *Myiarchus* flycatchers in Bolivia (Lanyon, 1978), *Henicorhina* wood-wrens in Ecuador (Halfwerk *et al.*, 2016), and *Ramphocelus* tanagers in Colombia (Sibley, 1958; Morales-Rozo *et al.*, 2017). The apparent paucity of hybridization between birds replacing each other with elevation in the Neotropics further suggests that elevational replacements did not originate through primary divergence in parapatry in the absence of barriers to gene flow, but instead via secondary contact of reproductively isolated populations. However, our inference of lack of hybridization between elevational replacements in the Andes is largely based on patterns of phenotypic variation; in most cases it remains to be seen whether genetic data reveal cryptic gene flow (Weir *et al.*, 2015).

While we cannot reject the hypothesis that closely allied species of *Scytalopus* replacing each other along the eastern slope of the Andes colonized such regions independently, our analyses indicate that parapatric speciation along the elevational gradient may have occurred there. Several of the species replacing each other with elevation in the Satipo Valley of Peru are closely related to each other, belonging to a clade of relatively recent origin in which mtDNA divergence is shallow and rates of speciation appear faster than in the rest of the genus (Cadena *et al.*, 2019b). Two members of this clade, *S. latrans* and *S. micropterus*, also replace each other with elevation in eastern Ecuador and Colombia. Because shallow divergence in putatively neutral loci and high rates of speciation may reflect rapid divergence mediated by adaptation in the face of gene flow, future studies should explicitly test predictions of ecological speciation (Smith *et al.*, 2005) to determine whether elevational replacements on the eastern Andean slope may indeed be uniquely explained by parapatric divergence. The same is true for the western slope of the northern Andes, where phylogeographic patterns suggest parapatric speciation may have occurred in amphibians and reptiles (Arteaga *et al.*, 2016; Guayasamin *et al.*, 2017).

Beyond examining patterns of relationships among species and populations, studies of the mechanisms underlying adaptation and of how adaptive evolution in the face of gene flow may lead to speciation along elevational gradients are needed (see Hua, 2016 for a theoretical perspective). In birds, for example, putatively adaptive variation with elevation has been documented in various traits influencing functions such as respiratory physiology (Scott, 2011; Dawson *et al.*, 2016; York *et al.*, 2017), thermoregulation (Scott *et al.*, 2008; Symonds & Tattersall, 2010), foraging (Kleindorfer *et al.*, 2006; McCormack & Smith, 2008), locomotion (Altshuler *et al.*, 2004; Milá *et al.*, 2009), and vocal signalling (Dingle *et al.*, 2008; Kirschel *et al.*, 2009). Whether any of such selective pressures may account for speciation by directly or pleiotropically influencing mating patterns in *Scytalopus* in the eastern Andean slope and in other groups is essentially unknown.

Given that elevational replacements more often reflect secondary contact than parapatric divergence, a central question involving the origin of non-overlapping ranges characterizing many species assemblages from tropical mountains remains unanswered. Do contrasting elevational distributions of species originate in allopatry or upon secondary contact? Diamond (1973) reasoned that elevational parapatry reflects the outcome of competitive interactions, whereby interspecific competition between formerly isolated species favors divergence of elevational ranges when they come into contact. Alternatively, contrasting elevational ranges may arise in allopatry, with the non-overlapping ranges of species one observes reflecting sorting processes, such that only species differing in elevational ranges *a priori* may successfully attain regional sympatry with segregation along mountain slopes (Cadena, 2007; McEntee *et al.*, 2018). A recent analysis revealed that sympatric sister species of birds in the tropics have more different elevational ranges than allopatric sister species regardless of their age, which was interpreted as evidence in favor of the hypothesis that elevational divergence is driven by competition upon secondary contact (Freeman, 2015). While abutting elevational ranges may indeed be maintained by competition in some cases (Cadena & Loiselle, 2007; Jankowski *et al.*, 2010; Freeman & Montgomery, 2016), other biotic and abiotic forces may also mediate species turnover with elevation (Elsen *et al.*, 2017). Moreover, because most cases of elevational replacements do not involve sister species, more work is necessary to determine the geographic context in which contrasting elevational ranges arise. Analyses incorporating species interactions into models of trait evolution (Nuismer & Harmon, 2015) while jointly considering the potential for such interactions to occur given geographic distributions of species (Drury *et al.*, 2016; Clarke *et al.*, 2017) would be a fruitful avenue for future studies on the topic (Quintero & Landis, 2019). Other mechanisms through which elevational distributions of species may change including tectonic processes of uplift or subsidence which may displace organisms vertically in passive fashion also merit consideration (Heads, 1989, 2005; Ribas *et al.*, 2007). *Scytalopus* tapaculos are well suited for additional studies on the dynamics of elevational ranges integrating ecology, evolution, and Earth history.

## Acknowledgments

We thank Ana Carnaval and Valentí Rull for inviting us to contribute this chapter. Andrés Cuervo, Niels Krabbe, Daniel Lane, Thomas Schulenberg, and V. Piacentini shared valuable information on the distributions of tapaculos. We are grateful to Ana Carnaval, Santiago Herrera, Ethan Linck, Ignacio Quintero, Glenn Seeholzer, Felipe Zapata, an anonymous reviewer, and CDC’s laboratory group for helpful discussion and comments on the manuscript. Fernando Ayerbe kindly allowed us to use his illustrations of birds in our figures. Paola Montoya, Gustavo Bravo and Elkin Tenorio provided measurement data, and Cooper French and Ethan Linck provided photographs for Figure 2. We received financial support from the Facultad de Ciencias at Universidad de los Andes (Programa de Investigación to CDC).

